# Functional cortico-subcortical reorganization after complete hemispheric disconnection for intractable epilepsy

**DOI:** 10.1101/707539

**Authors:** Thomas Blauwblomme, Athena Demertzi, Jean-Marc Tacchela, Ludovic Fillon, Marie Bourgeois, Emma Lositto, Monika Eisermann, Daniele Marinazzo, Federico Raimondo, Sarael Alcauter, Frederik Van De Steen, Nigel Colenbier, Steven Laureys, Volodia Dangouloff-Ros, Lionel Naccache, Nathalie Boddaert, Rima Nabbout

## Abstract

Hemispherotomy is a treatment for drug-resistant epilepsy with the whole hemisphere involved in seizure onset. As recovery mechanisms are still debated, we characterize functional reorganization with multimodal MRI in two children operated on the right hemisphere (RH). We found that interhemispheric functional connectivity was abolished in both patients. The healthy left hemispheres (LH) displayed focal hyperperfusion in motor and limbic areas, and preserved network-level organization. The disconnected RHs were hypoperfused despite sustained network-level organization. Functional connectivity was increased in the left thalamo-cortical loop and between the cerebelli. The classification probability of the RH corresponding to a minimally conscious state was smaller than for the LH. We conclude that after hemispherotomy, neurological rehabilitation is sustained by cortical disinhibition and reinforcement of connectivity driven by subcortical structures in the remaining hemisphere. Our results highlight the effect of vascularization on functional connectivity and raise inquiries about the conscious state of the isolated hemisphere.

## 1 Introduction

Epilepsy in children may involve an entire hemisphere due to developmental (malformations of cortical development, hemimegalencephaly), acquired (perinatal stroke) or progressive lesions (Rasmussen’s encephalitis, Sturge Weber Syndrome). Surgery is a first-line treatment option as drug resistance is common in these etiologies and because cognitive outcome is negatively impacted by the time of surgery. The surgical techniques have evolved from resection of the epileptic hemisphere (hemispherectomy) to disconnection procedures (hemispherotomy) where interhemispheric association bundles, projection and thalamo-cortical fibers are divided around the thalamus core, via a lateral (peri-sylvian) or midline approach (Baumgartner et al., 2017). This major surgery is associated with a 70% seizure freedom rate and a good functional outcome (Griessenauer et al., 2015; Ibrahim et al., 2015).

Despite living with only one hemisphere, operated children regain at least partial sensory motor function, do not worsen their cognitive skills, and may recover from language deficits, regardless of the operated side (Bulteau et al., 2015; Devlin et al., 2003). The understanding the reorganization of neural networks after hemispherotomy, however, suffers from methodological issues as these diseases are rare and imply young and often moderate to severely delayed children. The few available studies have focused on motor pathways probed by diverse medical imaging, and electrophysiological techniques, such as transcranial magnetic stimulation (Shimizu et al., 2000), somatosensory evoked potentials, positron emission tomography Müller et al. (1998), functional magnetic resonance imaging (fMRI) (Graveline et al., 1998; Holloway et al., 2000), tensor diffusion weighted imaging (Wakamoto et al., 2006), and combination of these techniques (Zhang et al., 2013). Collectively, these studies point to minimal motor plasticity changes in the remaining hemisphere, structural deteriorations in the affected hemisphere, and the ability to transfer motor and sensory functions from the resected hemisphere to the remaining one (Graveline et al., 1998; Hertz-Pannier et al., 2002).

A major question relates to the kind of cognitive processes still occurring in a fully disconnected hemisphere that lacks thalamo-cortical loops. Is this residual activity complex enough to enable a conscious or minimally conscious state (MCS, showing complex behavioral responses to external stimulation, such command following and pursuit of moving objects; Giacino et al., 2002) or does it rather correspond to non-conscious state such as observed in vegetative state/unresponsive wakefulness syndrome (VS/UWS, showing reflexive behaviors; Laureys et al. 2010)? We note that the here adopted hemipsherotomy model departs from the split-brain patients who typically show preserved thalamo-cortical activity in each of the two hemispheres that are disconnected (Gazzaniga, 2005), therefore it constitutes a unique opportunity to measure conscious functions in a more specific way.

Here, we characterize cerebral functional integrity and organization after complete hemispherotomy leading to seizure freedom in two children with Rasmussen’s encephalitis and hemimegalencephaly. By means of multimodal imaging, we aimed at exploring functional reorganization of both the remaining and the disconnected hemisphere at the cortical, subcortical and cerebellar level and at shedding light on the conscious capacities of the disconnected hemispheres.

## 2 Results

Both pediatric patients were studied postoperatively with perfusion and functional MRI. Patient 1 (MA) and patient 2 (JJ) were operated at 14 and 2 years of age respectively. The surgery involved the right hemisphere and aimed at complete hemispheric disconnection. A senior radiologist (NB) studied each patient’s T1 MRI images and confirmed the anatomical disconnection, including disruption of corpus callosum, anterior commissure, fornix, and internal capsule (***Figure 1***). After 5 years of follow up, both patients were seizure-free, with a good functional outcome (***Table 1***). Cerebral blood flow (CBF) and resting state fMRI functional connectivity measured whole-brain functional integrity, network-level organization, and subcortico-cortical interactions. Age-matched healthy controls for fMRI were obtained from the National Database for Autism Research (NDAR) (http://ndar.nih.gov). The dataset for MA included n=11 controls (1 female, mean age=16.8y±0.5SD, min=16, max=18). The dataset for JJ included n=9 controls (4 females, mean age=3.2y±0.3SD, min=3, max=4). ASL controls included 30 healthy subjects measured on the same MRI scanner in our institution (9 females, mean age = 10.3 ±3.2SD, min = 6, max = 18).

**Table 1.** Patient demographic characteristics

|  | MA | JJ |
| --- | --- | --- |
| Gender | male | male |
| Age at Sz onset | 13.6y | 10d |
| Sz semiology | Epilepsia Partialis Continua | Spasms |
| Age at surgery (yrs) | 14.3 | 2 |
| Age at MRI (yrs) | 17.6 | 4.6 |
| Preoperative EEG | PC central spikes | PC R temporooccipital spikes |
| Laterality | Right handed | NA |
| Etiology | Rasmussen encephalitis | Hemimegalencephaly |
| Seizure outcome | Engel score Ia, 5 years | Engel score Ia, 6 years |
| Motor outcome 1 | Hemi-paresis, -anopia, hypertonia | Hemi-paresis, -anopia, hypertonia |
| Motor outcome 2 | Walks, runs, eats, draws | Walks, runs, eats, draws |
| Functionality | Typical high school, 3 languages | Typical nursery school, bilingual |

Abbreviations: Sz= seizure, PC=pseudocontinuous

**Figure 1.**
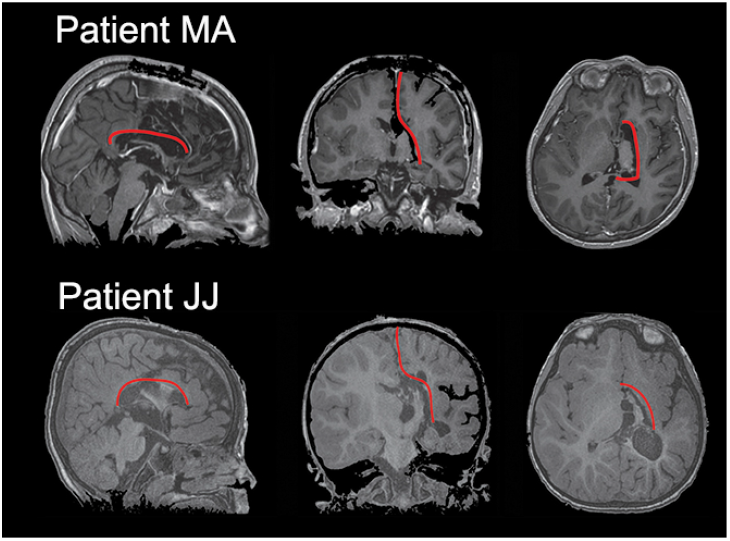
Surgical disconnection of the pathological right hemisphere in a case of Rasmussen’s encephalitis (MA) and hemimegalencephaly (JJ). The red line shows the surgical peri-thalamic disconnection after a midline approach. After an interhemispheric approach, complete callosotomy was performed, allowing access to the lateral ventricles. Perithalamic section of the white matter between the frontal and temporal horn disrupted the internal capsule, fimbria, anterior commissure, but left the major intra-hemispheric bundles (superior and inferior longitudinal fasciculi, uncinated fasciculus, cin-gulum, external capsule) untouched.

We first identified that interhemispheric functional connectivity was abolished. Pearson’s r correlations between each hemisphere’s gray matter averaged timeseries were just above zero for both MA (r= 0.09) and JJ (r= 0.03) who appeared as outliers among their controls (controls MA median: 0.50, min: 0.46, max: 0.78, 1st quartile: 0.48, 3rd quartile: 0.72; controls JJ median: 0.75, min: 0.48, max: 0.83, 1st quartile: 0.71, 3rd quartile: 0.81) (***Figure 2***)

**Figure 2.**
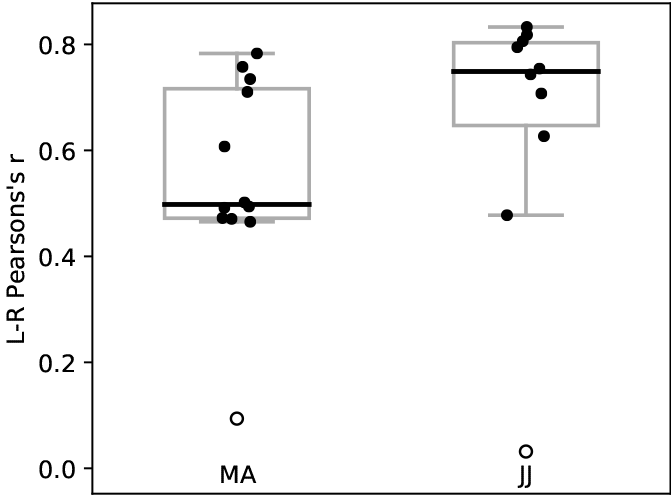
Interhemispheric connectivity was abolished in patients MA and JJ following complete hemispheric disconnection. Both patients showed correlation values just above zero and appeared as outliers (white circles) among their control subjects (MA controls, n=11; JJ controls, n=9). Boxplots represent median (thick line), interquartile range, minimum and maximum values.

Subcortically, the two patients showed increased connectivity between subcortical structures and the preserved left hemisphere. Thalamo-cortical connectivity increased ipsilaterally in the healthy left hemisphere in both cases (***Figure 3***) whereas the disconnected right hemisphere showed no residual thalamo-cortical functional connections. Both patients had increased connectivity between the two cerebelli, and between the healthy left cerebral hemisphere and both cere-bellar hemispheres (***Figure 3***). Patient JJ showed further right-sided ipsilateral cerebello-cortical connectivity, which was atypical given the surgical (***Figure 3*** yellow circle). This atypical pattern was mediated by the effect of the venous system. In particular, we found that the timeseries extracted from the superior sagittal sinus (SSS) predicted functional connections with the isolated right hemisphere in both patients. We noticed that the original cerebellar seed region was positioned over an area with functional connectivity predicted by the SSS. When a new seed was used which did not overlap with the SSS effect, no ipsilateral right connectivity was further observed (***Figure 3–1*** and ***Figure 3–2***).

**Figure 3.**
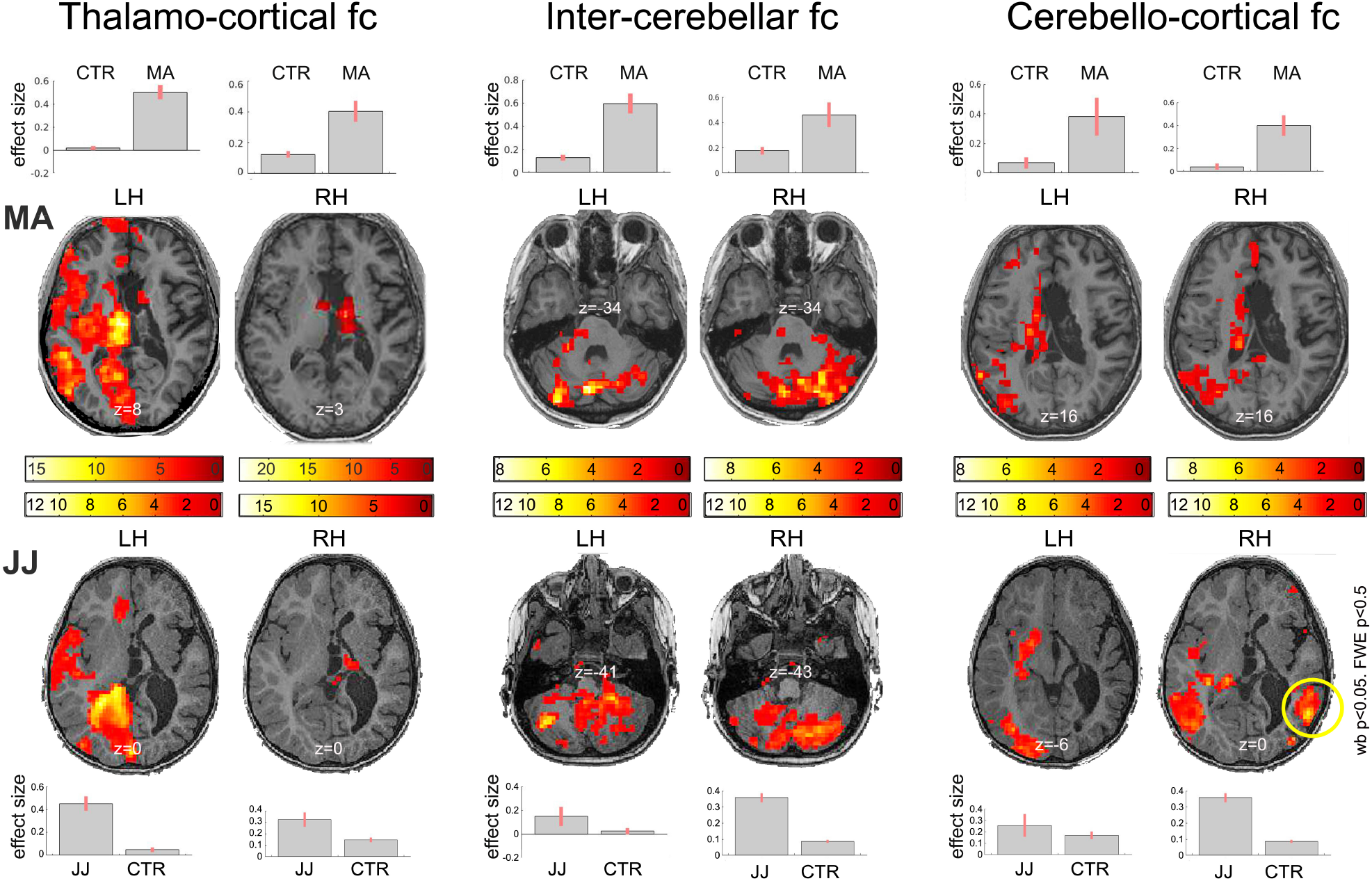
Mediation of subcortical structures in functional reorganization after complete hemispherotomy. Left: Thalamo-cortical functional connectivity (fc) increased ipsilaterally in the healthy left hemisphere in both patients, whereas the disconnected right hemisphere showed no residual thalamo-cortical connections. Middle: Both patients showed preserved and enhanced connectivity between the two cerebelli. Right Additionally, the two cerebelli had functional connections with the healthy left cerebral hemisphere but not with the isolated right. The atypical right-sided ipsilateral cerebello-cortical connectivity (yellow circle) seen in patient JJ was found to be an artifact mediated by the effect of the vascular system (superior sagittal sinus) and disappeared after regressing out that signal. Statistical maps are thresholded at whole-brain height threshold p<0.01, and cluster-level FWE p<0.05. Results are rendered on each patient’s normalized T1 image. Colorbars indicate t values. Bars indicate cluster-level contrast estimates (effect size) with 90% confidence intervals. Numbers in white refer to MNI slice coordinates (axial view)

The disconnected right hemisphere showed significant (p<0.05, FWE corrected) diffuse decreases in cerebral blood flow (CBF) values in both patients compared to healthy individuals (MA, mean= 16.7 mL/100mg/mn ±15.4SD; JJ, mean= 34.1±17.3SD; controls mean= 45 mL/100mg/mn ±2.7SD; FWE corrected p=0.05; ***Figure 4A***). Crossed cerebellar hypoperfusion was also noted in both patients. After regressing out the SSS effect, intrinsic fMRI functional connectivity in the disconnected right hemi-spheres showed lateralized network-level organization in large-scale (default mode, frontoparietal, salience) and sensory systems (auditory, motor, visual). Seeds ROIs placed on the right hemi-sphere showed no contralateral connectivity in either patient (whole brain p<0.01, cluster-level FWE p<0.05) (***Figure 4B***), in contrast to the bi-lateral connectivity typically observed in healthy controls (***Figure 4–1*** and ***Figure 4–2***).

**Figure 4.**
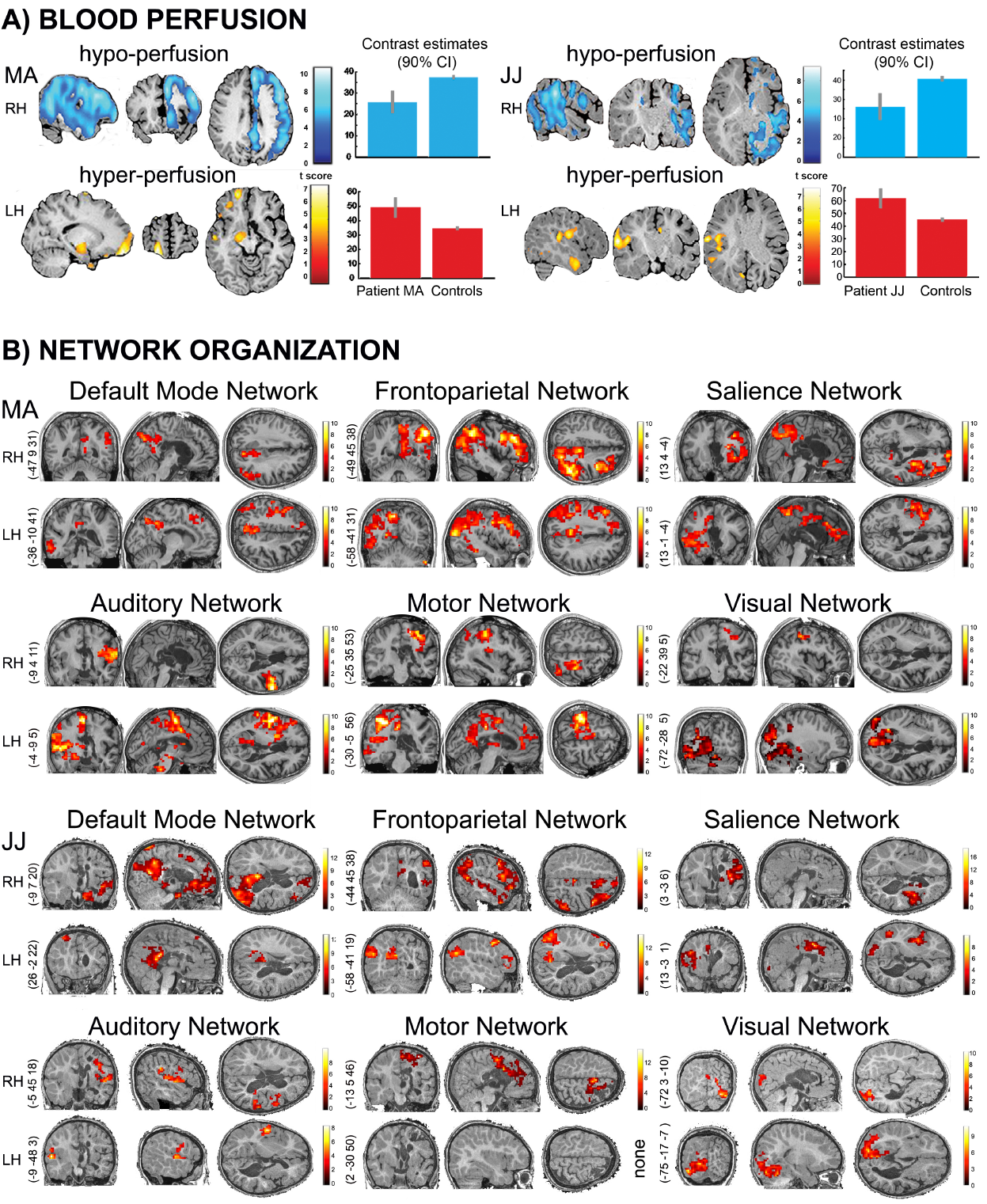
Whole-brain functional organization was laterized after complete hemispherotomy. A) In terms of blood perfusion, cerebral blood flow was lower within the disconnected right hemisphere (blue regions, FWE p<0.05) compared to healthy controls both for patient MA and JJ. At the same time, focal hyperperfusion was observed in the remaining left hemisphere in both patients (blue regions, FWE p<0.05) compared to healthy controls. Bars represent averaged contrast estimates across the identified cluster with 90 % confidence interval (whiskers). The statistical maps are rendered on the patient’s normalized T1 image. B) In terms of functional connectivity, both patients showed network organization in six representative systems. The connectivity appeared lateralized within the right (RH) and left hemisphere (LH) and did not show contralateral connectivity transfer. Of note is the preserved yet restricted network-level connectivity in the isolated right hemisphere even after the regression of the vascularization effect of the superior sagittal sinus. Statistical maps are thresholded at whole-brain height threshold p<0.01, and cluster-level FWE p<0.05. Results are rendered on each patient’s normalized T1 image. Colorbars indicate t values. Side numbers refer to MNI slice coordinates.

The healthy left hemisphere showed localized significant (p<0.05, FWE corrected) increases in CBF values in both patients compared to healthy individuals (MA, mean= 66.2 mL/100mg/mn ±10.5SD; JJ, mean= 69.05 mL/100mg/mn±11.6SD, controls mean= 47 mL/100mg/mn ±1.8SD). Hyperperfusion was located in the motor operculum, amygdala, temporal and frontal pole in MA and in the temporal pole and sensorimotor operculum in JJ (***Figure 4A***). After regressing out the SSS effect, intrinsic fMRI functional connectivity showed lateralized network-level organization in large-scale (default mode, frontoparietal, salience) and sensory systems (auditory, motor, visual). Seeds ROIs placed on the left hemisphere showed no contralateral connectivity in either patient (whole brain p<0.01, cluster-level FWE p<0.05) (***Figure 4B***) in contrast to the bilateral connectivity typically observed healthy controls (***Figure 4–1*** and ***Figure 4–2***).

To evaluate the residual cognitive processes occurring in the right disconnected hemispheres, we aimed at testing how they would be classified by an algorithm we previously designed to distinguish cortical activity of VS/UWS versus MCS patients. More specifically, we adapted this algorithm to classify separately functional connectivity of each hemisphere (intra-hemispheric functional connectivity). Two key features per hemisphere for each patient were classified among patients who demonstrated disorders of consciousness. We previously identified that connectivity in bilateral temporal cortices and in occipital regions separated 20/22 patients with disorders of consciousness according to clinical behavioral assessments, suggesting that intrinsic brain connectivity can be used as a means to provide reliable information about the level of consciousness in a data-driven way (Demertzi et al., 2015). Here, by considering each hemisphere separately for the classification, the features were separated in half resulting in temporal and occipital regions for each hemisphere. The analysis included patient MA only as he was the most comparable to the included subjects in the training set. A linear support vector machine classifier was trained on 26 patients in MCS and 19 patients in VS/UWS and generalized on patient MA. For the isolated right hemisphere, the probability of belonging to the class of MCS was 0.65. For the preserved left hemisphere, the probability of belonging to the class of MCS was 0.96 (***Figure 5***).

**Figure 5.**
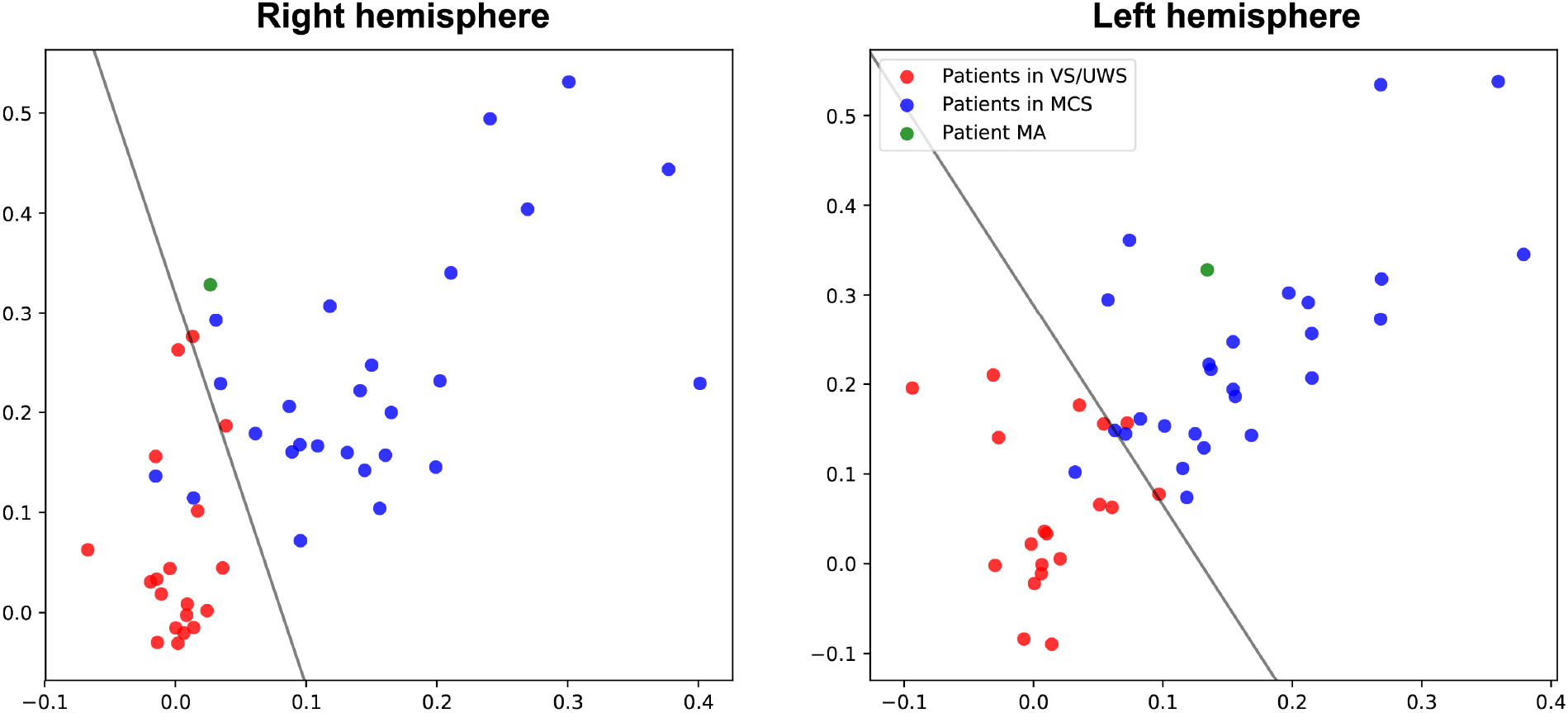
The contribution of each hemisphere to the state of consciousness. For patient MA (green) the isolated right hemisphere was closer to the class of patients in vegetative state/unresponsive wakefulness syndrome (VS/UWS, in red; showing reflexive behaviors) and had low chances to be classified among patients in minimally conscious state (MCS, in blue; showing complex behaviours to external stimulations but who remain unable to communicate). At the same time, the preserved left hemisphere was classified toward the class of MCS with higher probability. The line represents the decision boundary between the two classes as estimated by the linear support vector classifier.

## 3 Discussion

We studied brain organization after right hemispherotomy in two pediatric patients treated for intractable epilepsy. By means of cerebral hemodynamics, we tested whether intrinsic architecture provides insights about cortical and subcortical reorganization after complete hemispheric disconnection. The effects of this procedure were evident on both ASL and resting state fMRI analysis, which respectively showed rCBF and functional connectivity patterns clearly lateralized. Specifically, we found widespread reduction of the cerebral blood flow along with preserved intrinsic functional connectivity in the disconnected hemisphere, whereas contralaterally cerebral blood flow and connectivity increased in the remaining cortex and subcortical structures.

In the disconnected right hemisphere, we identified reductions in cerebral blood flow both ipsilaterally and with the contralateral cerebellum. Reduced hemispheric metabolic demands have been already described as secondary to thalamic stroke and after thalamotomy for tremor, correlating with cognitive outcome (Baron et al., 1992, 1986). More recently, perfusion studies with ASL highlighted crossed cerebellar and ipsilateral diaschisis (i.e., decreases in blood flow and metabolism in areas unaffected by the lesion) with reduced perfusion of rCBF after thalamic or putaminal hemorrhage (Noguchi et al., 2015). Such significant decreases in neuronal metabolism may be due to reduced neuronal activity secondary to the loss of afferent inputs (Gold and Lauritzen, 2002). For example in rodents, cerebellar cortex interneurons and Purkinje cells decreased their firing rate after inducing focal cerebral ischemia (Gold and Lauritzen, 2002). Similarly for hemispherotomy, section of the perithalamic white matter tracts suppresses seizure expression due to the interruption of the cortical projection bundles to the brainstem and spinal cord. In the meantime, there is interruption of the ascending reticular activating system (an essential polysynaptic pathway for arousal through increases of cortical excitability; Steriade 1996). Therefore, the isolated right hemisphere can be said that it is characterized by low neuronal activity after hemispheric disconnection.

In the remaining left hemisphere, we identified CBF increases notably in the somatosensory cortex, and mesial/lateral temporal region compatible with elevated neuronal activity. These results are in line with a previous TMS study after hemispherotomy which showed that motor cortex excitability was enhanced after the surgery (Shimizu et al., 2000). Following the lesion paradigm of stroke, up regulation of contralateral homotopical areas has already been reported in the left middle cerebral artery territory, where aphasic patients showed an up-regulation of the right Broca-homologue region during language tasks in the sub-acute phase (Saur et al., 2006). Such a phenomenon could rely on the disinhibition of the healthy hemisphere. An in vivo imaging study indeed showed that after selective unilateral stroke there was stroke-induced enhancement of responses (evoked by stimulation of the limb contralateral to the stroke in the spared hemisphere), with subcortical connections (rather than transcallosal projections) mediating the novel activation patterns (Mohajerani et al., 2011). After hemispherotomy, this disinhibition could be related to the callosotomy despite that most of callosal fibers transmit excitatory glutamatergic inputs. Indeed, in rodents after contralateral sensory stimulation of the somatosensory cortex, the firing of layer 5 pyramidal neurons was inhibited when paired with ipsilateral stimulation, suggestive of interhemispheric inhibition (Palmer et al., 2012). The localization of the focal CBF increases is further reminiscent of activation tasks in fMRI and neurophysiological studies after hemispherotomy, pointing to residual function originated from the remaining healthy hemisphere in expected networks. As such, MEG somatosensory evoked potentials elicited ipsilateral responses in the primary somatosensory cortex in three patients with residual sensory function after hemispherectomy (Yao et al., 2013). Combined neurophysiological and fMRI showed ipsilateral activation of the sensory motor region during passive movement of the hand in a location similar to the movements of the other hand, yet with a greater spatial extent (Holloway et al., 2000). Interestingly, fMRI investigating motor ankle dorsiflexion before and after rehabilitation showed an extended motor network after rehabilitation with an overlap of ipsi- and contralateral regions of interest in the supplementary motor area and primary sensorimotor cortices (de Bode et al., 2007) The corticospinal tracts is the sole motor output pathway after hemispherectomy and do not undergo obvious modifications after surgery at the supra spinal levels as assessed by DTI studies (Wakamoto et al., 2006). At the level of the spinal cord, the formation of ipsilateral descending pathways following neonatal hemidecortication can be due to a loss of balance in cortical activity between the two hemispheres (Umeda and Funakoshi, 2014). The same pattern of reorganization was also previosuly noted in the language network, where pre- and post-operative language task activation fMRI after left hemispherotomy showed a putative transfer of the network in the right hemisphere (Hertz-Pannier et al., 2002; Liégeois et al., 2008). Taken together, the preserved left hemisphere increased perfusion might reflect a mechanism of disinhibited neural activity after the callosotomy.

We also found that patients showed increased subcortical connectivity in their remaining left hemisphere. More precisely, we identified increased thalamo-cortical connectivity, which could be expected as the thalamus is a major node in brain networks (Hwang et al., 2017). This postoperative connectivity pattern was similar to previous a report (Ibrahim et al., 2015). Therefore, we can postulate that high-order thalamic nuclei may be key drivers for functional adaptation in the non-disconnected hemisphere. This hypothesis is supported by electrophysiological recordings in non-human primate and rodents. In macaques, the pulvinar mediates synchrony between the V4 area and temporo-occipital cortex in the alpha band during attentional visual task, facilitating the transmission of information about attentional priorities in distinct non-contiguous modules of the visual cortex (Saalmann et al., 2012). Also in rodents during attention control tasks, prefrontal cortex neurons spike along with dorsomedial thalamic neurons, that supervise their connectivity. Optogenic suppression or activation of thalamic neurons could modulate the results of the cognitive tasks through inhibition or recruitment of further prefrontal cortex neurons by the thalamic neurons, thus demonstrating that thalamic high-order relay can control cerebral cortex connectivity (Schmitt et al., 2017).

Cerebellar connectivity was also modified in both patients as compared to the control groups. Crossed cerebello-thalamo-cortical connectivity was enhanced, in agreement with previous imaging studies after cerebral cortex extended lesions. Indeed, higher cerebellar cortical glucose utilization was noted in PET studies in children younger than one year of age along with decreased benzodiazepine receptor binding in the dentate nucleus contralateral to the lesion (Niimura et al., 1999). These findings are related to anatomical modifications that were noted both in animal models and after hemispherectomy, with expansion of the afferent and efferent fibers of the cortico-ponto-cerebello-rubro-thalamic system (Govindan et al., 2013; Olmstead et al., 1983). As the cerebellum is highly involved in learning (Medina and Lisberger, 2008) and as it modulates cerebral excitability via its thalamic inhibition (Jayaram et al., 2011; Spampinato and Celnik, 2017) we hypothesize that after hemispherotomy increased connectivity between cerebellar hemispheres consequently shapes cortical reorganization. Moreover, its role in motor recovery may also involve bilateral spinal efferences through the Ruber nuclei to compensate the loss of the cortico-spinal tract as already demonstrated in animal models and after stroke in humans (Rüber et al., 2012; Siegel et al., 2015).

Interestingly, we further observed that patient JJ initially showed right-sided ipsilateral cerebello-cortical connectivity. When we further investigated this atypical result to our hypothesis, we found that there was a mediation of the effect of the vascular system. Indeed, when the signal of the SSS was regressed out during signal denoising in the preprocessing phase, this ipsilateral connectivity disappeared. The effect of large blood vessels refers to systemic low frequency oscillations (sLFO 0.1Hz), usually present in vascularized tissue (e.g. fingertips, toes). Such sLFOs are non-neuronal signals which are also included in the BOLD fMRI variance and their origin are cardiac, respiratory, and peripheral (Tong et al., 2013). The sLFO were used to track cerebral blood flow by determining the time-lags between these signals and the BOLD signals from different voxels (Tong and Frederick, 2014; Tong et al., 2016). For example, it has been shown that sLFO signals in carotid arteries can be used to estimate the time delays which precede signals found in the brain (Tong et al., 2016). In the future, we expect that more refined identification of the venous system after hemispherotomies will shed more light on the neuronal and non-neuronal connectivity of the disconnected hemispheres.

The reason why intrinsic network functional connectivity was preserved in brain regions which do not contribute to behavioral output remains an open question. A working hypothesis is that this activity is critical for the development of synaptic connections and maintenance of synaptic homeostasis at large (Pizoli et al., 2011) which is reduced, yet preserved, in covert or unconscious conditions (Demertzi et al., 2019; Heine et al., 2012). The disconnected right hemisphere might therefore be a model of unilateral disorder of consciousness (Bruno et al., 2011). For instance, global CBF decreases were noted before in patients in MCS (Liu et al., 2011). In the cohort of MCS patients, along with patients in VS/UWS, there were further reductions in connectivity strength within large-scale and sensory networks when compared to healthy individuals (Demertzi et al., 2014, 2015; Vanhaudenhuyse et al., 2010). Such observations raise queries as to the role of such intrinsic organization in the absence of behavioral merit. Here, using a previously developed classifier for discriminating states of consciousness after severe brain injury, we found that for patient MA the classification prediction of his left hemisphere was high for the MCS class, hence indicating a higher consciousness state. At the same time, his right hemisphere was mostly identified among connectivity values seen in unresponsive/vegetative state patients. This finding was in line with our hypothesis that the isolated hemisphere might contribute in conscious state less compared to the preserved left, and that thalamo-cortical processing plays a necessary role for conscious processing. Also, note that MCS has been recently reinterpreted as a Cortically Mediated State (Naccache, 2018), indicative of a class of behaviors revealing the active contribution of cortical networks, rather than a univocal conscious state. Under such hypothesis, MCS does not relate to consciousness but to a necessary but insufficient condition for conscious processing. The fact that a disconnected hemisphere could be classified as MCS is therefore not univocal and future studies should precise its cognitive and conscious/unconscious status. The classification of the level of consciousness, then, should be interpreted mostly as indicative information rather than an absolute model for consciousness function. This is because the used classifier was formed on a qualitatively different population to separate the state of consciousness, namely adult patients suffering severe brain damage. Considering, though, the sparsity in the number of patients having received complete hemispherotomy, we think that even such coarse testing sheds light on the ongoing debates about the role of the isolated hemisphere in behavior and paves the way for more thorough examination of the preserved capacities of isolated brain tissue by more interventional means.

In conclusion, hemispheric disconnection is a major neurosurgical procedure for epileptic children with hemispheric epilepsy, leaving one hemisphere mostly functional. After complete hemispherotomy, whole-brain functional organization appears laterized, allowing us to postulate that in the healthy hemisphere cortical disinhibition and enhanced connectivity, driven by subcortical structures through preexisting networks, mediate neurological recovery. Our results point to methodological considerations concerning the effect of the vascular system in functional connectivity after hemispherotomy and raise further inquiries about the cognitive and conscious state of the isolated hemisphere.

## 4 Methods and Materials

### 4.1 Subjects and surgical technique

In a series of 10 hemispherotomy cases performed between 2013 and 2014, eight patients accepted to be enrolled in the CREIM imaging protocol approved by the local ethics committee. Six patients had to be excluded from the study due to motion artifacts during the fMRI acquisition, ongoing seizure activity in the disconnected hemisphere, or because of behavioral problems precluding full fMRI protocol without sedation. Preoperative MRI was not performed because none of the patients could do perform it without sedation.

Both children were operated by a midline vertical hemispherotomy (Baumgartner et al., 2017). After an interhemispheric approach, complete callosotomy was performed, allowing access to the lateral ventricles. Peri-thalamic section of the white matter, between the frontal and temporal horn, disrupted the internal capsule, fimbria, anterior commissure, but left the major intra hemispheric bundles (superior and inferior longitudinal fasciculi, uncinated fasciculus, cingulum, external capsule) untouched (***Figure 1***).

For the fMRI analysis healthy controls were included as a reference in the second-level analysis of the performed one sample t-tests. These healthy subjects were age-matched and were obtained from the National Database for Autism Research (NDAR) (http://ndar.nih.gov) scanned on 3T scanners (Siemens Magnetom TrioTim or General Electric SignaHDxt). The dataset for patient MA included n=11 controls (1 female, mean age=16.8y±0.5SD, min=16, max=18). The dataset for patient JJ included n=9 controls (4 females, mean age=3.2y±0.3SD, min=3, max=4). For the ASL analysis, 30 healthy pediatric controls were used as previously reported (Boisgontier et al., 2018).

### 4.2 Data Acquisition

MRI scanning was performed without sedation 39 months and 31 months after surgery for MA and JJ respectively.

For the ASL data, the 3D ASL sequences were acquired on a GE Signa HDxt 1.5T system (General Electric Medical System, Milwaukee, USA) using a twelve-channel head-neck-spine coil including morphological sequences (3D T1-weighted images, Axial T2 FLAIR, Diffusion) non-contrast perfusion imaging with 3D pseudo continuous ASL MRI (pcASL). The acquisition included 80 axial partitions (field of view 240 × 240 × 4 mm3; acquisition matrix 8 spiral arms in each 3D partition, 512 points per arm; TE 10.5 ms; TR 4428 ms; Post Labeling Delay 1025 ms; flip angle 155°; acquisition time 4 min 17 s). For the fMRI session, data were acquired on a GE Discovery MR750 3T system and included 300 functional MRI T2*-weighted images acquired with a gradient-echo echo-planar imaging (EPI) sequence using transverse slice orientation and covering the whole brain (39 slices, slice thickness = 3mm, repetition time = 2000ms, echo time = 34ms, voxel size = 3.125×3.125mm, flip angle = 90°). A structural T1 magnetization prepared rapid gradient echo sequence (120 slices, repetition time = 2300ms, echo time = 2.47ms, voxel size = 1.0×1.0×1.2mm, flip angle = 9°).

### 4.3 Data Analysis: fMRI

#### 4.3.1 FMRI preprocessing

Preprocessing was performed using SPM12 (https://www.fil.ion.ucl.ac.uk/spm/software/spm12/) as implemented in Matlab (Mathworks Inc., Sherborn, MA, USA), including slice-time correction, realignment, segmentation of structural data, normalization of functional and structural data into standard stereotactic MNI space and spatial smoothing using a Gaussian kernel of 6mm full-width at half-maximum. For functional data, the three initial volumes were discarded to avoid T1 saturation effects. Motion artifact detection and rejection was performed with the artifact detection toolbox (ART toolbox, https://www.nitrc.org/projects/artifact_detect/).

An image was defined as an outlier or artifact image if the head displacement in x, y, or z direction was greater than 2mm from the previous frame, or if the rotational displacement was greater than 0.02 radians from the previous frame, or if the global mean intensity in the image was greater than 3 standard deviations from the mean image intensity for the entire resting scan. Outliers in the global mean signal intensity and motion were subsequently included as nuisance regressors within the first-level general linear model so that the temporal structure of the data would not be disrupted. For noise reduction, the anatomical component-based noise correction (aCompCor) method (Behzadi et al., 2007) was utilized as implemented in the CONN functional connectivity toolbox (v.16b) (Whitfield-Gabrieli and Nieto-Castanon, 2012). This approach models the influence of noise as a voxel-specific linear combination of multiple empirically estimated noise sources by deriving principal components from noise regions of interest (ROIs) and by including them as nuisance parameters within the general linear models. Specifically, the anatomical image for each subject was segmented into white matter (WM), gray matter (GM), and cerebrospinal fluid masks (CSF). For patient MA, the default mask as provided in CONN was used. For patient JJ, due to his developing brain morphology, a template of a 2-year old was used. The template was constructed with longitudinal and cross-sectional group-wise registrations of a set of longitudinal images acquired from 95 typical infants. For the sake of the current analysis, the 2yr old group-wise anatomical (intensity) model with skull was used (Shi et al., 2011). To minimize partial voluming with GM, the WM and CSF, masks were eroded by one voxel which resulted in smaller masks than the original segmentations. The eroded WM and CSF masks were then used as noise ROIs: timeseries were extracted from the unsmoothed functional volumes to avoid additional risk of contaminating WM and CSF signals with gray matter signals. Five principal components of the signals from WM and CSF noise ROIs were removed with regression. Residual head motion parameters (three rotation and three translation parameters, and six parameters representing their first-order temporal derivatives) were further regressed out. A temporal high-pass filter [0.008Hz inf] was used for estimating the interhemispheric correlations. A temporal band-pass filter [0.008-0.09Hz] was used for estimating functional connectivity as classically performed in resting state analyses.

To account for potentially detected functional connectivity in the disconnected hemisphere, we opted to isolate the effect of vascularization on the BOLD neuronal signals. Neuronal low frequency activity changes should not be confused with systemic low frequency oscillations (sLFO 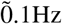) usually present in vascularized tissue (e.g. fingertips, toes). sLFO are non-neuronal signals which are also included in the BOLD fMRI variance and their origin are cardiac, respiratory, and peripheral Tong et al. (2013). The sLFO, especially in large veins such as the superior sagittal sinus (SSS), were shown to highly correlate with the fMRI global signal, namely the averaged BOLD signal across the brain whitech reflects primarily capillary and venous blood (Funnell et al., 2000; Honey et al., 2009). Using the 3Dslicer (v4.8.1 r26813, 18) (Kikinis et al., 2014) on the patient’s T1/T2 or FLAIR raw data, we identified the superior sagittal sinus manually. The identified segments were then coregistered and normalized on MNI space to match the dimensions of the normalized functional images. The extracted timeseries from the SSS were then used as a noise ROI to regress out the SSS effect. The functional connectivity associated with the SSS signal is summarized in ***Figure 3–1*** and ***Figure 3–2***.

### 4.4 Interhemispheric correlations

Interhemispheric correlations were estimated using Pearson’s r between each subject’s right and left gray matter averaged timeseries. In particular, the segmented smoothed gray matter images were separated between the left and the right hemisphere using the fslroi function. These half-hemisphere gray matter images were used as masks to extract the averaged timeseries from each subject’s denoised data. Pearson’s r correlation coefficient was then calculated on these averaged values.

### 4.5 Functional connectivity

Functional connectivity adopted a seed-based correlation approach. Seed-correlation analysis uses extracted blood oxygenation level-dependent timeseries from a region of interest (the seed) and determines the temporal correlation between this signal and the timeseries from all other brain voxels. Seed-based brain parcellation followed each patient’s anatomical constraints. They were 5mm-radius sphere ROIs referring to pertinent intrinsic connectivity networks, such as the default mode, frontoparietal, salience, motor, auditory, and visual (Demertzi et al., 2015; Raichle, 2011; Smith et al., 2009) (Supplementary Tables 1 and 2 in ***Appendix A***). Due to the young age of patient JJ, the ROI for left thalamus with connections to DMN followed the coordinates from our previous work (Alcauter et al., 2014). The averaged timeseries were used to estimate whole-brain correlation r maps that were then converted to normally distributed Fisher’s z transformed correlation maps to allow for group-level comparisons. One-sample t-tests were ordered to estimate network-level functional connectivity for patients MA and JJ separately using their corresponding control subjects as a reference group [modelling 1(patient) 0 (controls)]. To allow for visualization of potential contralateral connectivity, results were considered significant a liberal whole-brain height threshold p<0.01, with cluster-level corrections for multiple comparisons at FWE rate p<0.05.

### 4.6 Classification of consciousness level

The assessment of consciousness level was tested by means of a modified version of a previously developed classifier targeting to separate patients with diverse consciousness states (Demertzi et al., 2015). The classifier was based on a connectivity pattern including three regions/features, including bilateral superior temporal/precentral gyri and occipital areas. This pattern came from the difference in connectivity between a group of patients in a minimally conscious state (showing complex behavioral responses to external stimulation, such command following and pursuit of moving objects) and patients in vegetative state/unresponsive wakefulness syndrome (showing reflexive behaviors). Here, this pattern was normalized on each patient’s normalized anatomical image and was separated in half to represent the right and left hemisphere. This led to a modified classification scheme with values/features coming from two clusters per hemisphere per patient (i.e. temporal and occipital regions, left and right). These features were extracted as follows. First, whole-brain connectivity was estimated using six seed regions per patient. For MA, these seeds refer to the R precentral gyrus (58 −6 11), R superior temporal gyrus (44 −6 11), L precentral gyrus (−53 −6 8) L superior temporal gyrus (−44 −6 11), L occipital cortex (−6 −83 43), R occipital cortex (6 −83 43). Then, by means of the REX toolbox (www.nitrc.org/projects/rex/), we used these connectivity maps as Sources and the half brain masks as ROIs to extract cluster-level averaged connectivity values (average z-values across the two clusters) for the left and right hemisphere separately, leading to two features per hemisphere.

We used a linear support vector machine classifier, trained on 26 patients in MCS (21 males; mean age=46 years; 13 traumatic, 13 non-traumatic of which 3 were anoxic; 20 patients assessed >41 month post-insult), and 19 patients in VS/ UWS (12 males; mean age; 1 traumatic, 18 non-traumatic of which 11 anoxic; 13 patients assessed 41 month post-insult). The discrimination performance was summarised with the area under the curve (AUC) calculated from the receiver operator characteristic (ROC) curve. For a binary classification system, the ROC pits the detection probability, commonly referred to as sensitivity, against the probability of false alarm (1 - sensitivity). These probabilities are empirically estimated by moving the decision cut-off along the sorted values of a continuous variable and by evaluating its relation to the true label. The AUC can then be conveniently used to summarise the performance, where a score of 0.5 is uninformative and equals to random guessing whereas a score of 1 amounts to perfect classification and 0 to total confusion, indicating negative correlation between the score and the label. The probability in belonging to the class of MCS was estimated by fitting the distribution of the samples with regards to the optimal linear combination of features (w) (Platt, 2000). A sigmoid function was fitted from the distributions of the signed distances separating the train samples and w. This sigmoid fit was eventually used to monotonically transform the signed distance separating the test samples and w into a meaningful probability.

### 4.7 Data Analysis: Arterial Spin Labelling

#### 4.7.1 ASL preprocessing

Due to the poor spatial resolution of the ASL images, the registration was done in an indirect but robust way including the preprocessing of structural T1 data (Blauwblomme et al., 2016). The preprocessing steps were achieved using the Voxel Based Morphometry toolbox (Ashburner and Friston, 2000) (VBM8) (http://www.neuro.uni-jena.de/vbm), implemented in Matlab (Mathworks Inc., Sherborn, MA, USA). First, T1 and ASL data are converted from DICOM to NIFTI format. Then, native T1 images are segmented into gray matter, white matter and cerebrospinal fluid classes. For some patients, when the segmentation failed in the operated hemisphere due to large defects in the white matter, it was necessary to fill the removed area with a “simulated white matter signal” (corresponding to a gaussian distribution with the same mean and standard deviation intensities than the white matter in the contralateral hemisphere) and segment this new image with VBM8. With the gray matter and white matter segmentation images, a brain mask was built to extract the brain from the native T1 image followed by normalization on a 7 year-old-brain atlas obtained with Template-O-Matic Toolbox TOM8 (http://www.neuro.uni-jena.de/software/tom/) as implemented in SPM8 (Wilke et al., 2008). ASL images were co-registered on the native gray matter image to take into account the potential movement of patient during T1 and ASL acquisition including translation and rotation. The coregistered ASL was then normalized using the deformation field obtained during the T1 normalization process. Eventually the normalized ASL images are smoothed using a 10mm isotropic filter. Voxel-based analyses were performed on smoothed and normalized ASL images within a grey matter mask in under SPM8, as previously (Blauwblomme et al., 2016). At the individual level, voxel-based analysis was performed using the general linear model, comparing the patient to a control group of 30 healthy pediatric controls according to a methodology previously described (Boisgontier et al., 2018). Results were interpreted with a significance level set at whole-brain p=0.05 FWE, and p=0.001 non-FWE corrected.

## 5 Acknowledgments

AD is a Research associate at the Fund for Scientific Research (FNRS), Belgium. SL is a Research Director at the Fund for Scientific Research (FNRS), Belgium.

## A APPENDIX 1

**Table 2.**
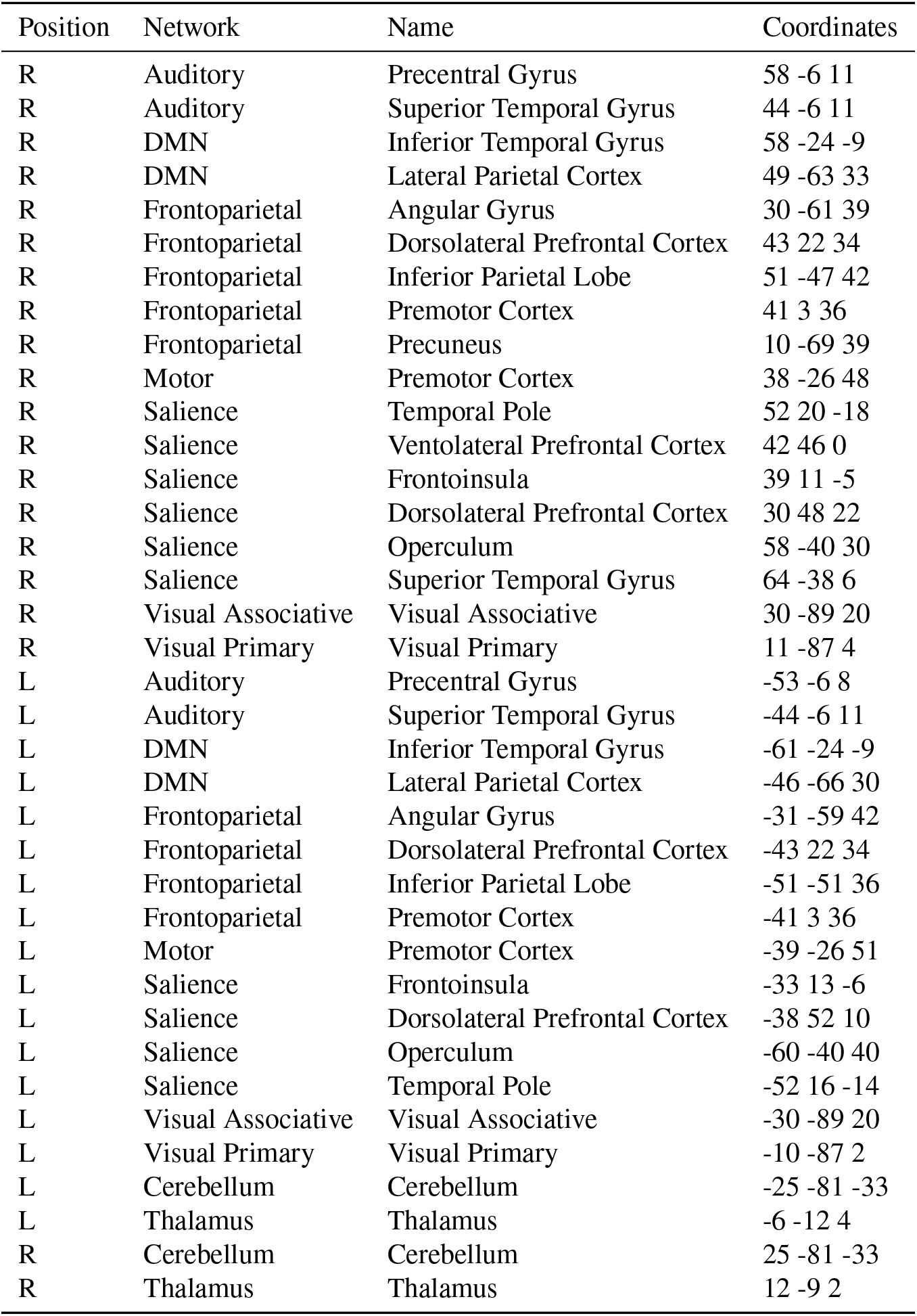
Regions of interest for patient MA. Regions of interest were spheres (5mm-radius) designed based on each patient’s anatomical constraints referring to the most pertinent intrinsic connectivity networks.

**Table 3.**
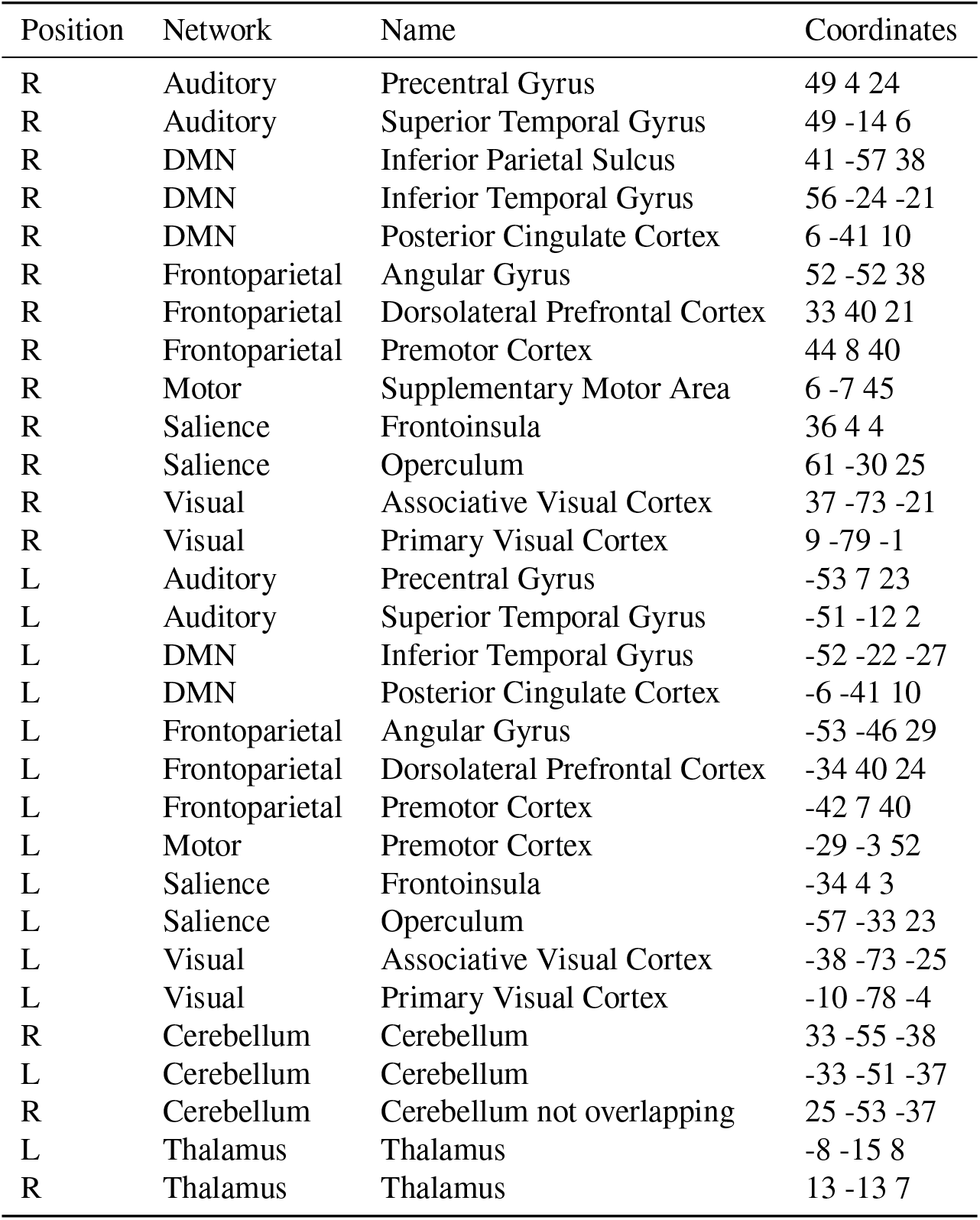
Network-level organization of regions of interest (patient JJ). Regions of interest were spheres (5mm-radius) designed based on each patient’s anatomical constraints referring to the most pertinent intrinsic connectivity networks.

**Figure 3–Figure supplement 1.**
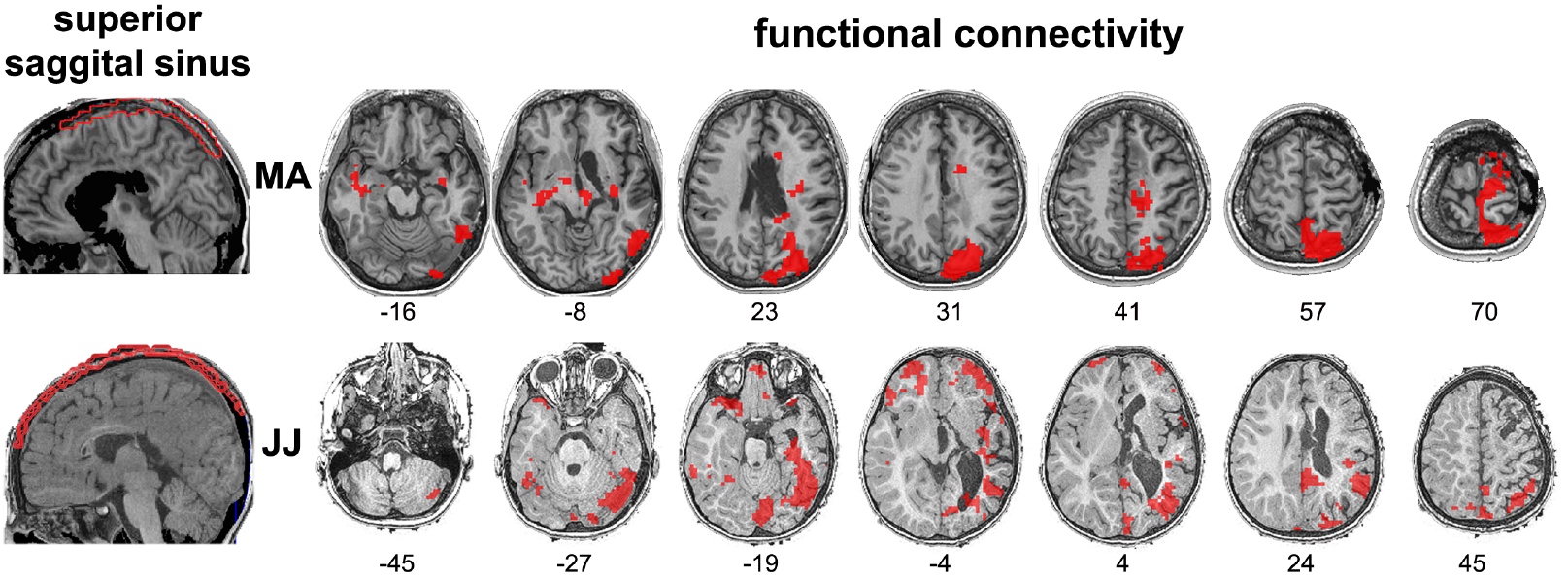
Explaining the cerebello-cortical functional connectivity of the pathological hemisphere. The vascularization system was hypothesized as the main source of inter-hemispheric transfer of cortico-subscortical functional connectivity because it is the only physiological system shared by the two hemispheres. Considering the superior saggital sinus (SSS) as a seed region, functional connectivity was predicted bilaterally. Statistical maps are thresholded at whole-brain height p<0.01, FWE cluster level p<0.05 and rendered on each patient’s normalized MRI.

**Figure 3–Figure supplement 2.**
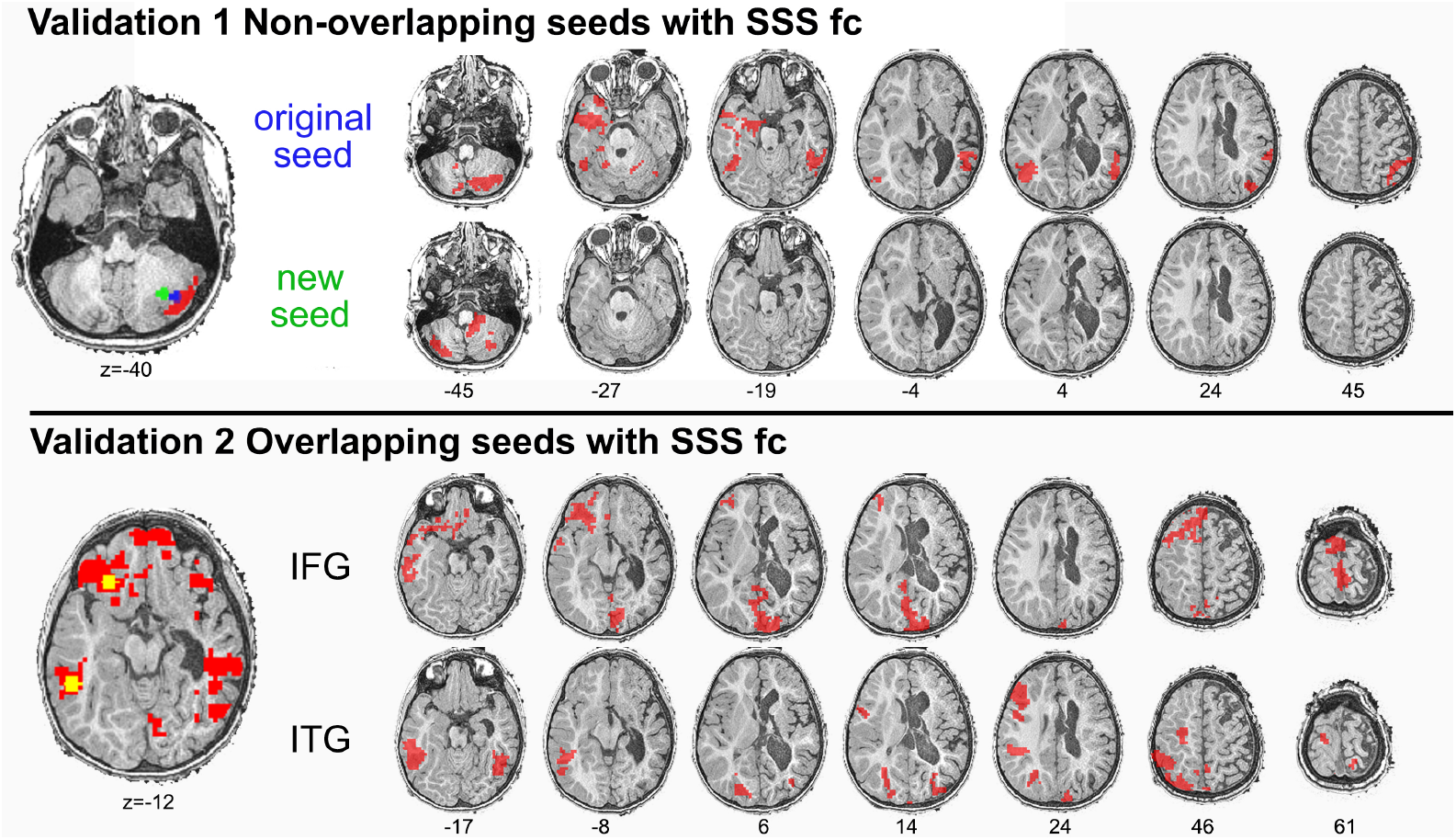
To verify this effect on patient JJ, two validation tests were performed. First, we noted that original cerebellar seed (blue) was positioned over an area with functional connectivity (fc) predicted by the SSS timeseries (red region). When a new cerebellar seed was used (green), which was not overlapping with the SSS effect, no ipsilateral right connectivity was observed. Second, when two other seeds (yellow, over the inferior frontal gyrus-IFG and the inferior temporal gyrus-ITG) were placed on regions which were functionally connected with the SSS timeseries, both ipsi- and contralateral connectivity was predicted, verifying the confounding effect of the vascularization of the SSS on the isolated right hemisphere. Statistical maps are thresholded at whole-brain height p<0.01, FWE cluster level p<0.05 and rendered on the patient’s normalized MRI.

**Figure 4–Figure supplement 1.**
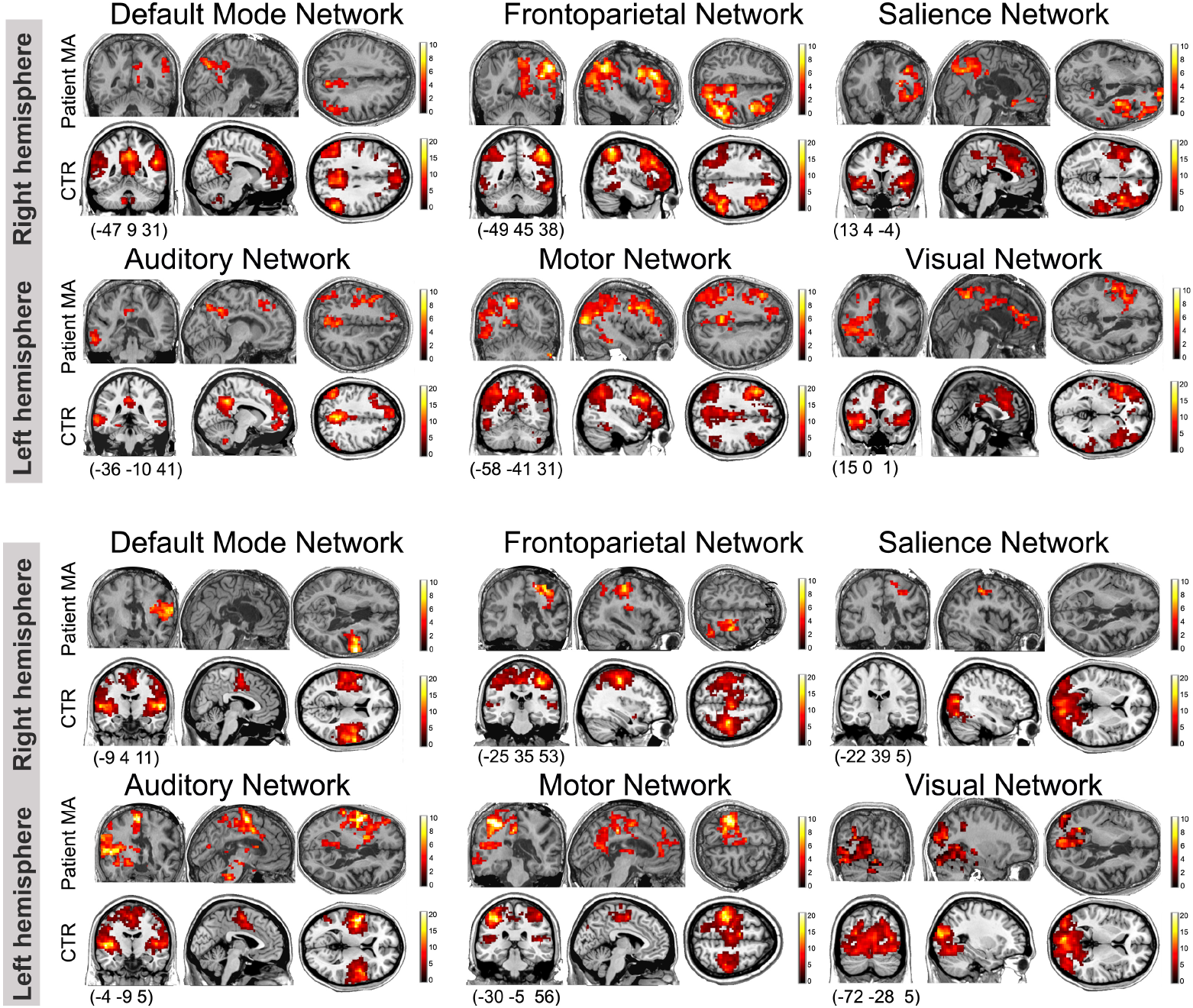
Network-level organization of intrinsic functional connectivity in the right and left hemisphere as estimated in patient MA and his healthy control subjects (CTR, n=11). Of note is the bilateral connectivity in the control group for right- and left-sided regions of interest. This interhemispheric connectivity effect was not observed in patient MA, who showed lateralized connectivity restricted to each hemisphere in the six studied networks. Statistical maps are thresholded at whole-brain height threshold p<0.01 and at FWE p<0.05 (cluster-level correction) and rendered on the patient’s normalized T1 image and on a stereotaxic template for his controls (MRIcron, ch2 template), with coronal, sagittal and axial views. Colorbars indicate t values. Bottom numbers refer to MNI slice coordinates.

**Figure 4–Figure supplement 2.**
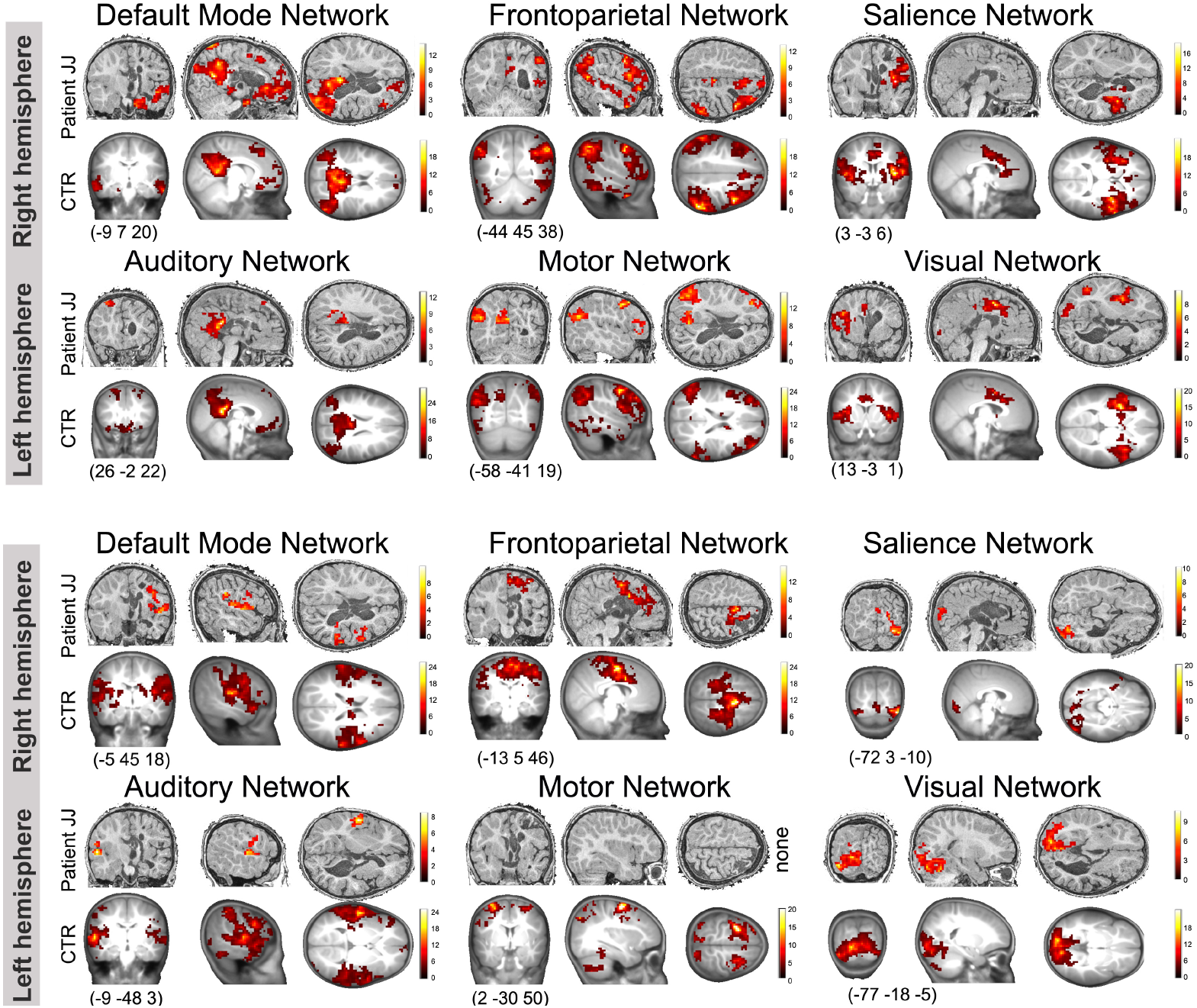
Network-level organization of intrinsic functional connectivity in the right and left hemisphere as estimated in patient JJ and his healthy control subjects (CTR, n=9). Of note is the bilateral connectivity in the control group for right- and left-sided regions of interest. This interhemispheric connectivity effect was not observed in patient JJ, who showed lateralized connectivity restricted to each hemisphere for the six studied networks. Statistical maps are thresholded at whole-brain height threshold p<0.01 and at FWE p<0.05 (cluster-level correction) and rendered on the patient’s normalized T1 image and on a normalized 2 year old infant template for his controls, with coronal, sagittal and axial views. Colorbars indicate t values. Bottom numbers refer to MNI slice coordinates.

